# Chromatin Network Retards Droplet Coalescence

**DOI:** 10.1101/2021.03.02.433564

**Authors:** Yifeng Qi, Bin Zhang

## Abstract

Nuclear bodies are membraneless condensates that may form via liquid-liquid phase separation. The viscoelastic chromatin network could impact their stability and may hold the key for understanding experimental observations that defy predictions of classical theories. However, quantitative studies on the role of the chromatin network in phase separation have remained challenging. Using a diploid human genome model parameterized with chromosome conformation capture (Hi-C) data, we studied the thermodynamics and kinetics of droplet formation inside the nucleus. Dynamical simulations predicted the formation of multiple droplets for protein particles that experience specific interactions with nucleolus-associated domains (NADs). Coarsening dynamics, surface tension, and coalescence kinetics of the simulated droplets are all in quantitative agreements with experimental measurements for nucleoli. Free energy calculations further supported that a two-droplet state, which is often observed for nucleoli seen in somatic cells, is metastable and separated from the single-droplet state with an entropic barrier. Our study suggests that protein-chromatin interactions facilitate the nucleation of droplets, but hinders their coarsening due to the correlated motion between droplets and the chromatin network: as droplets coalesce, the chromatin network becomes increasingly constrained. Therefore, protein-chromatin interactions arrest phase separation in multi-droplet states and may drive the variation of nuclear body numbers across cell types.

## INTRODUCTION

Nuclear bodies are pervasive in eukaryotic cells and play a diverse set of functions [1], including RNA metabolism, transcriptional regulation [2], genome organization [3] etc. They are membraneless structures that mainly consist of protein and RNA molecules [4]. Their lack of a lipid-rich barrier allows rapid exchange of components with the nucleoplasm in responses to environmental cues and stress signaling. Nuclear bodies also effectively increase the local concentration of enzymes involved in particular functions to facilitate more efficient cellular reactions [5].

Increasing evidence supports that nuclear bodies function as biomolecular condensates formed via liquid-liquid phase separation (LLPS) [4, 6–8]. They exhibit round morphologies and dynamic fluid properties [9]. Two nuclear bodies can fuse into larger condensates following coalescence dynamics consistent with simple liquids [10]. In addition, their assembly was shown to be concentration-dependent, and the coarsening and growth dynamics can be quantitatively modeled with classical theories of phase separation [11]. At the molecular level, detailed mechanistic models for LLPS are beginning to emerge as well. In particular, low complexity domains and intrinsically disordered regions are enriched in many of the proteins associated with nuclear bodies [12–16]. These features enable non-specific, multivalent interactions that drive the formation of dynamical condensates.

However, several aspects of nuclear bodies appear to defy predictions from classical nucleation and phase separation theories. In particular, the stable equilibrium state shall correspond to a single condensate that minimizes the surface energy [17]. On the other hand, multiple nucleoli (~2-5) are commonly observed in somatic cells [18], and the number can increase to 10^2^ in *Xenopus laevis* oocyte [10]. Quantitative experiments enabled with optogenetic approaches further revealed an anomalous exponent that deviates from classical predictions for the coarsening dynamics in living cells [19].

Nuclear bodies are known to be in close contact with chromatin via interactions mediated by long non-coding RNAs or chromatin-binding proteins [20]. For example, nucleoli are sites for ribosome biogenesis, and ribosomal DNA (rDNA) can be seen inside and adjacent to nucleoli [21]. Chromosome segments other than the nucleolus organizer regions (NORs) that enclose rDNA were found to associate with nucleoli as well and are collectively noted as nucleolus-association domains (NADs) [8, 22]. In addition, promyelocytic leukemia protein (PML) bodies [23], speckles [24, 25] and Cajal bodies [26] are all in spatial proximity with chromatin. Since chromatin forms a viscoelastic network spanning the nucleus, its interactions with nuclear bodies could impact the thermodynamics and kinetics of phase separation. Such interactions might hold the key to understanding the abnormal behaviors mentioned above.

We carried out molecular dynamics simulations to investigate LLPS in the nucleus with a computational model that includes protein particles and the chromatin network. We represented the chromatin network using a diploid human genome model that provides explicit polymer configurations for individual chromosomes. Interactions within and among chromosomes were optimized based on chromosome conformation capture (Hi-C) experiments to ensure in vivo relevance. The simulated dynamical process of droplet growth, coarsening, and coalescence are in quantitative agreement with experimental measurements. Importantly, our simulations predicted the formation of multiple droplets, much like the coexistence of several nucleoli seen in the nucleus. We showed that a two-droplet state is metastable and separated from the single-droplet state with an entropic barrier. The barrier arises from the chromatin network, which becomes more constrained upon droplet coalescence. Protein-chromatin interactions correlate the motion between the chromatin network and the droplets, and stronger interactions were shown to produce more droplets. Our study provides insight into the critical role of the chromatin network on nuclear body formation.

## RESULTS

### Phase Separation with Chromatin Network Leads to Multiple Droplets

Leveraging a recently introduced computational model for the diploid human genome [27, 28], we studied the impact of the chromatin network on phase separation inside the nucleus. We modeled the genome at the one megabase (Mb) resolution as a collection of 46 chromosomes inside a spherical confinement (Fig. 1). Each chromosome is represented as a string of beads, which can be assigned as one of three types, *A*, *B*, or *C*. *A* and *B* correspond to the two compartment types that contribute to the checkerboard patterns typically seen in Hi-C contact maps [29, 30], and *C* marks centromeric regions. Interactions among the beads were optimized to reproduce various average contact probabilities determined from Hi-C experiments using the maximum entropy optimization algorithm [31, 32] (see Supplementary Materials).

**FIG. 1.**
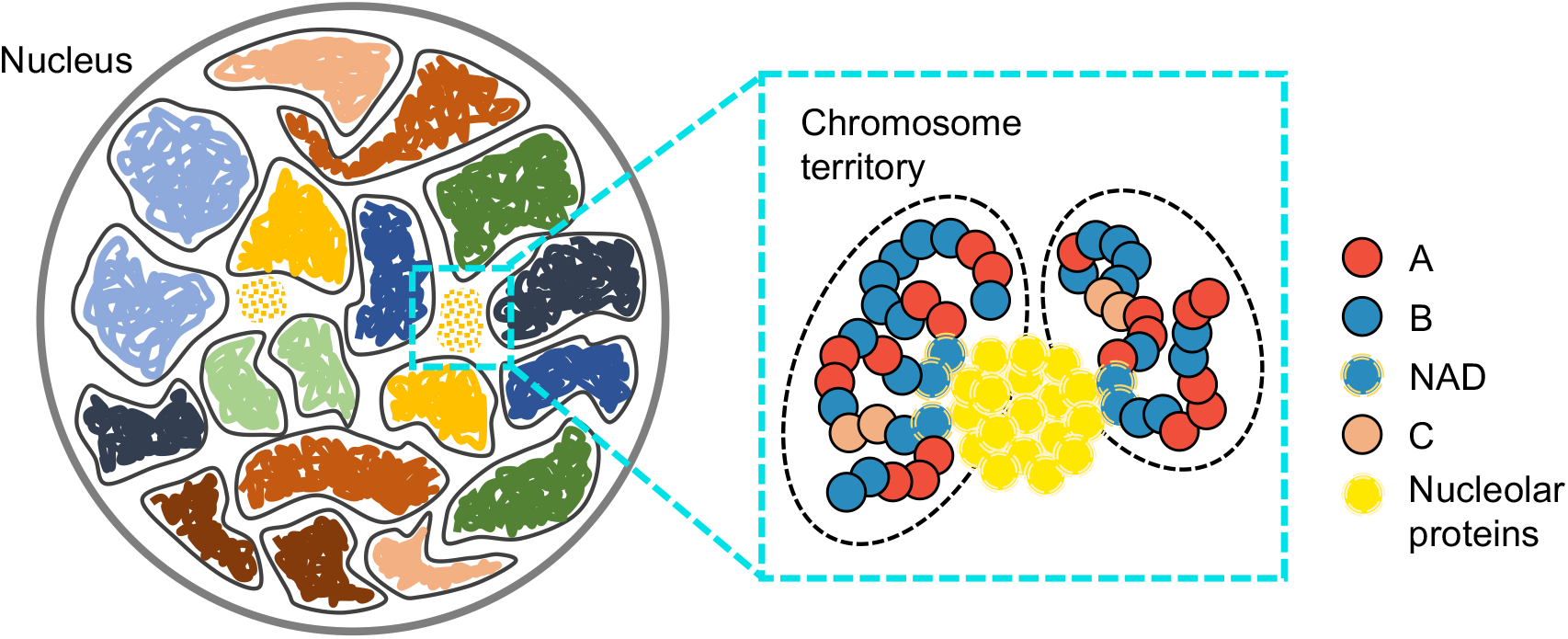
Overview of the computational model used for studying phase separation inside the nucleus. The diploid human genome model represents each one of the 46 chromosomes as a string of beads confined in the nuclear envelope. Each bead is identified as compartment type *A*, *B*, or centromeric region *C*. Protein particles share favorable interactions to promote phase separation and bind specifically with nucleolus associated domains (NADs).

We introduced additional coarse-grained particles to model protein molecules. Without loss of generality, we focused on nucleolar proteins that are key components of nucleoli [8]. These particles share favorable interactions with each other and with nucleolus-associated domains (NADs), which are chromatin regions strongly associated with nucleoli [33]. A non-specific interaction term was also introduced between protein particles and non-NAD chromatin regions. The Lennard-Jones potential was used for protein-protein and protein-chromatin interactions, with the interaction strength chosen as *ϵ* = 1.8 *k*_B_*T* and 1.0 *k*_B_*T* for specific and non-specific interactions, respectively. *k*_B_ is the Boltzmann constant, and *T* is the temperature. The strength of the specific interactions was found to stabilize droplets that share comparable surface tension as that of nucleoli (see Supplementary Materials). The total number of protein particles and their size were chosen based on the concentration of nucleolar proteins and the volume fraction of nucleoli (see Supplementary Materials).

Starting from an equilibrated genome structure and randomly distributed protein particles (Fig. 2A), we carried out a total of twelve independent molecular dynamics simulations. In all but one case, the protein particles aggregated into multiple droplets that persisted to the end of the simulations (Fig. 2B). This result contrasts with simulations performed without the chromatin network, where proteins will always aggregate into a single droplet (Fig. S1). Consistent with observations from fluorescence recovery after photobleaching (FRAP) experiments [35, 36], these droplets exhibit liquid-like property. As shown in Fig. S2, protein particles forming the droplets undergo dynamic exchange with surrounding nucleoplasm while maintaining the droplet size (~ 1 *μ*m) on the ten-of-minutes timescale. The droplets are preferentially localized at the interior of the nucleus (Fig. S3A). Because of their close association with these droplets, NADs are closer to the nuclear center than other heterochromatin regions as well [37]. However, not all NADs bind to the droplets, and a significant fraction of them localize towards the nuclear envelope (Figs. S3B-C). Two classes of NADs that vary in nuclear localization have indeed been observed in prior studies [38].

**FIG. 2.**
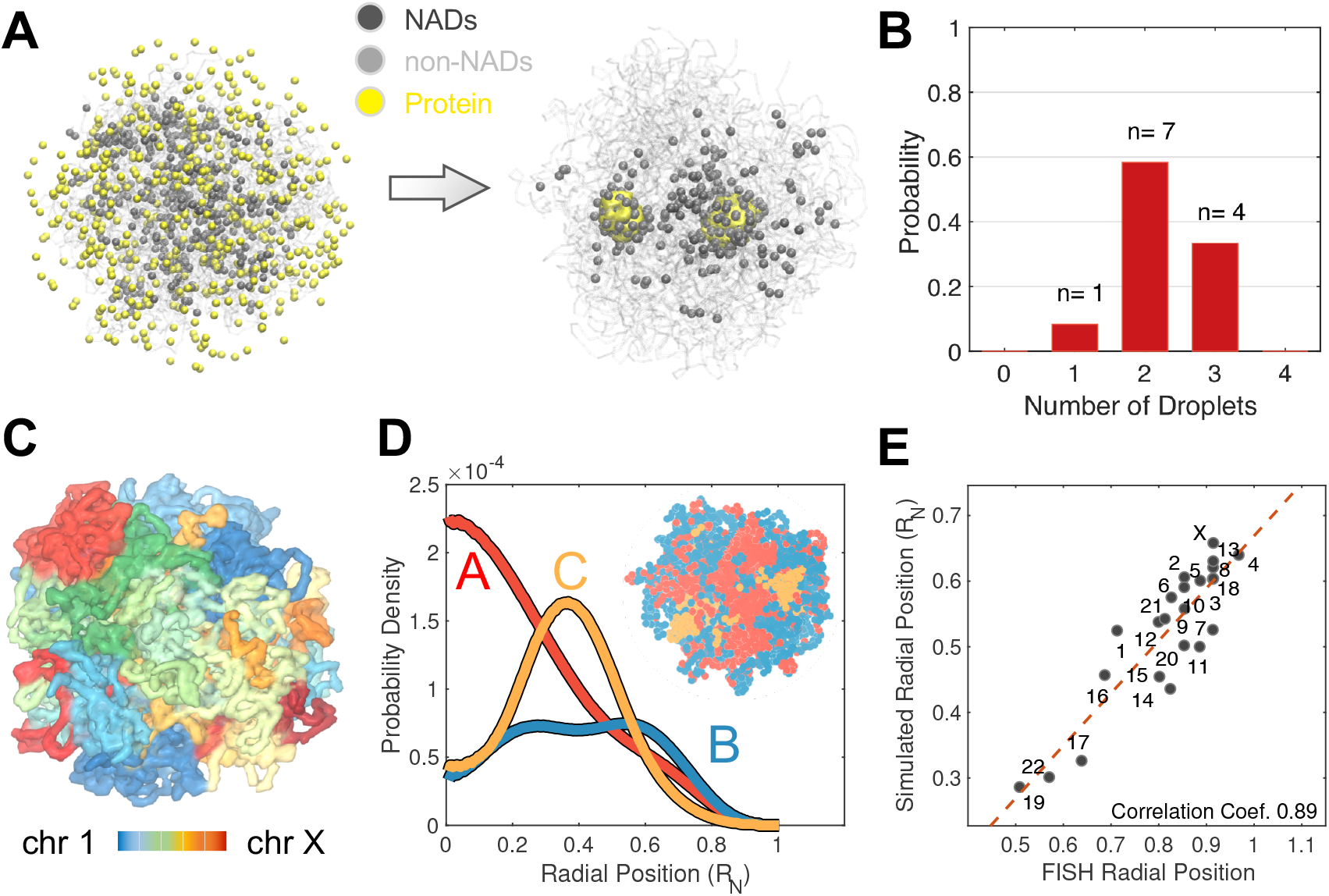
Multiple droplets form in dynamical simulations of the nucleus model. (A) Representative initial (left) and final (right) configurations obtained from simulations, with protein particles in yellow, NADs in black, and the rest of the genome in grey. (B) Probability distribution of the number of droplets observed at the end of simulation trajectories. (C) Representative configuration of the genome that illustrates the formation of chromosome territories. (D) Radial distributions of the different chromatin types that support their phase separation and preferential nuclear localization. An example genome configuration is shown as the inset, with the three types colored in red (*A*), blue (*B*), and orange (*C*), respectively. (E) Simulated radial chromosome positions correlate strongly with experimental values [34]. Homologous chromosomes were averaged together. *R*_N_ is the radius of the nucleus used in polymer simulations.

Analyzing the simulated genome structures, we found that global features of the genome organization are preserved after introducing protein particles, including the formation of chromosome territories [39–41], the compartmentalization of heterochromatin/euchromatin [42–44], and the clustering of centromeric regions [45–47], as shown in Figs. 2C-D. The simulated chromosome radial positions agree well with experimental values [34], and the Pearsons correlation coefficient between the two is 0.89 (Fig. 2E).

### An Entropic Barrier Hinders Droplet Coalescence

The emergence of a long-lived state with multiple droplets, while in contrast with predictions of the classical nucleation theory [48], is consistent with experimental observations [49–52]. As discussed in the *Introduction*, it is common to observe two nucleoli in normal cell types, and the number increases significantly in cancer cells [53, 54]. To better understand the stability of the multi-droplet state, we computed the free energy profile as a function of the radius of gyration (*R_g_*) of protein particles forming the droplets. As the size of individual droplets remains stable, changes in *R_g_* will be mainly driven by variations in the distance of the two droplets. However, different from a simple center-of-mass distance, which becomes ill-defined if the lists of protein particles participating in droplet formation are not updated on the fly, *R_g_* is relatively invariant with respect to the flux of particles between the two droplets (Fig. S4). Umbrella sampling and temperature replica exchange were used to enhance conformational exploration.

As shown in Fig. 3A, the free energy profile exhibits two basins. While the basin at *R_g_* ≈ 6.0 corresponds to the two droplet state observed before, an additional minimum with all protein particles participating in a single droplet appears at smaller values (≈ 3.0). The two basins are separated from each other with a transition state at *R_g_* ≈ 4.5. Representative configurations at the transition state show a dumbbell shape with the establishment of a thin bridge between the two droplets. Consistent with predictions of the classical nucleation theory, the one droplet state remains as the global minimum. However, the merging of the droplets is kinetically constrained due to the presence of a barrier that is approximately 10 *k*_B_*T* in height.

**FIG. 3.**
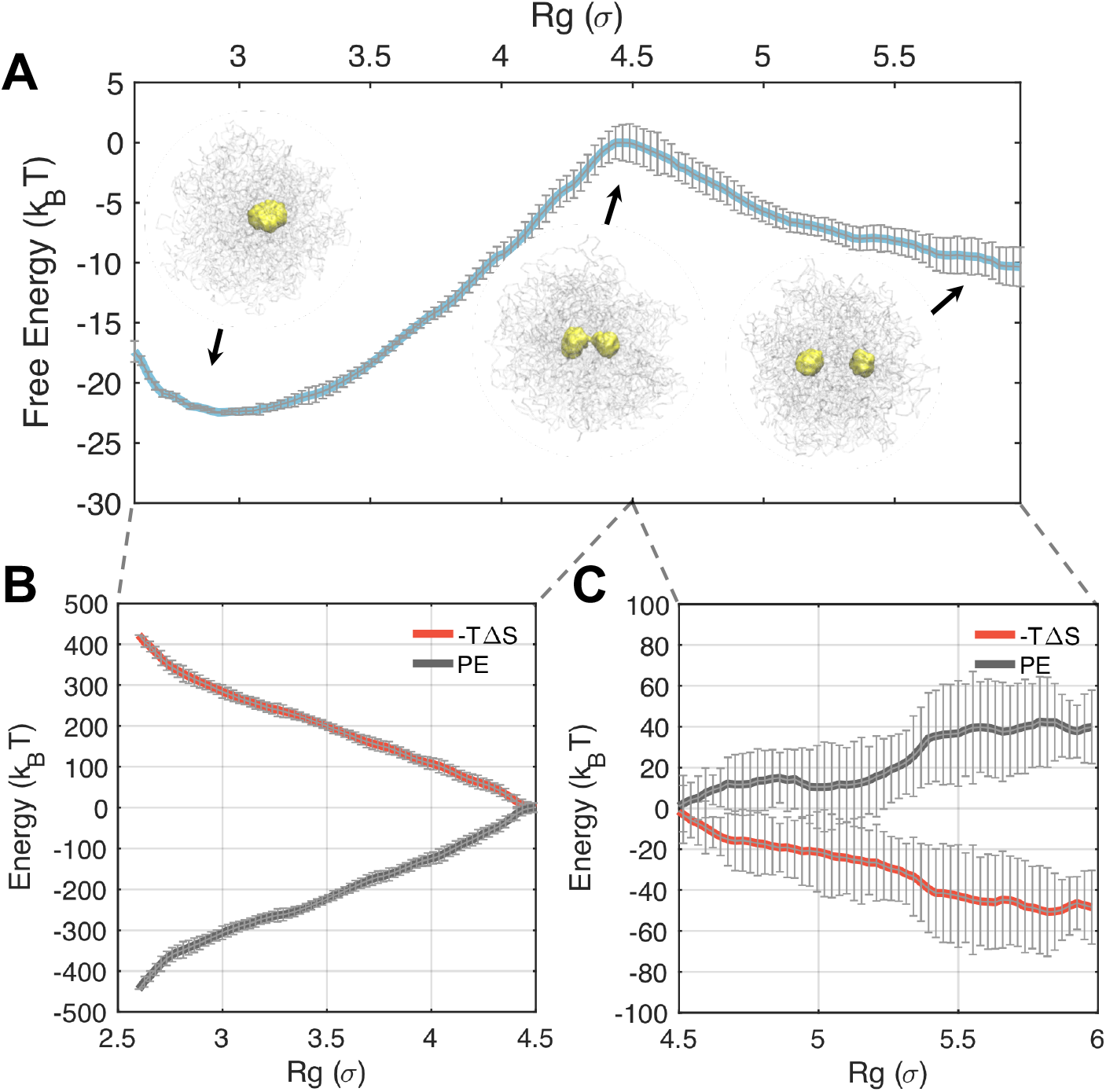
An entropic barrier hinders droplet coalescence and drives the metastability of a two-droplet state. (A) Free energy profile as a function of the radius of gyration (*R_g_*) that effectively measures the distance between two droplets. (B,C) Energetic (black) and Entropic (red) contributions to the free energy profile before (C) and after (B) the coalescence barrier. Error bars were calculated as standard deviation of the mean.

To reveal the nature of the barrier, we decomposed the free energy into entropic and energetic contributions. Using the free energy profiles at different temperatures (Fig. S5), we computed the entropy change along the collective variable with finite difference methods [55, 56]. As shown in Figs. 3B-C, contributions from the entropy (−*T* Δ*S*) continue to increase as *R_g_* decreases, and the droplets coalesce. While restricting the motion of the two droplets to smaller distances is naturally unfavorable, the entropic penalty is intensified here due to the increasingly restricted motion of chromosomes as well. As mentioned before, protein particles form extensive contacts with NADs via specific interactions, and such contacts enforce correlative motions between droplets and chromosomes. The potential energy, on the other hand, favors droplet merging and decreases continuously along the collective variable due to the increase in protein-protein and protein-chromatin contacts. Therefore, the transition state arises due to the presence of the chromatin network, and dissolving the polymeric topology of chromosomes indeed removes the barrier (Fig. S6).

While *R_g_* suffices for monitoring the progression of droplet coalescence, whether it serves as a “good” reaction coordinate or not requires further investigation. In particular, a good reaction coordinate should provide insight into the bottleneck that limits the reaction. Furthermore, trajectories initialized from the identified transition state will have an equal probability of committing to the reactant or product, or the so called committor probability adopts a value of 0.5 [57, 58]. Otherwise, the transition state may bear little relevance to the reaction, and mechanisms derived from it can be misleading. To evaluate the significance of the transition state for droplet coalescence, we carried out additional simulations starting from random configurations with *R_g_* values at ~ 4.5. For each configuration, we initialized ten independent 200,000-step-long simulations with randomized velocities. We then counted the number of simulations that end up with the two-droplet state versus the single-droplet state. As shown in Fig. 4, among all these simulations, 56% of them led to the single-droplet state while the rest 44% ended up in the two-droplet state. These results strongly support the usefulness of *R_g_* for studying coalescence and the relevance of the identified transition state for mechanistic interpretation. Since the chromatin network was not included for defining the reaction coordinate and can vary significantly at a given value for *R_g_* (Fig. S7), it may play a secondary or passive role in mediating coalescence.

**FIG. 4.**
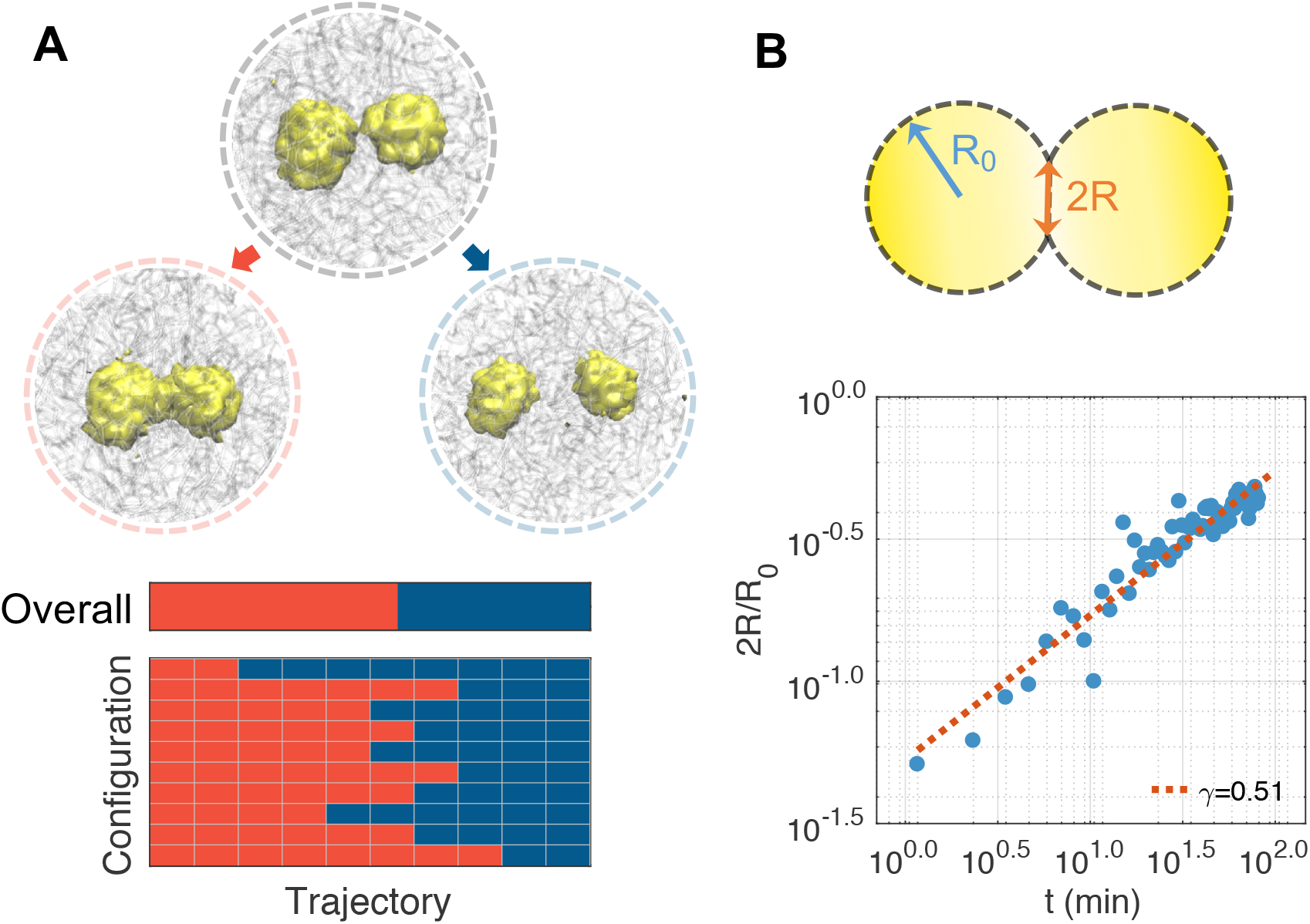
Dynamical characterization of droplet coalescence. (A) Trajectories initialized from transition state configurations share approximately equal probability of committing to the two and single droplet state. The number of trajectories that ended up in the single droplet state is shown as red blocks in the bottom panel. Example configurations of the single-droplet (left), transition (middle), and two-droplet (right) state are shown on top. (B) Time evolution of the radius of the neck region between two droplets.

We further isolated trajectories initialized from the transition state that led to the single-droplet state and computed the evolution of the neck radius as a function of time. As shown in Fig. S8, the neck region was identified as the minimum of the density profile of protein particles along the principal axis with the largest variance. As shown in Fig. 4B, by plotting the normalized neck radius (2*R*(*t*)*/R*_0_) with respect to the time, we obtained a power-law relationship with exponent 0.51. *R*_0_ is the average radius of the droplets before fusion. This exponent agrees with the experimental value determined for nucleoli [52] and suggests that droplet coalescence proceeds in the low Reynolds number regime dominated by viscous effects from the outer fluid, i.e., the nucleoplasm [59, 60].

### Chromatin Network Gives Rise to Slow Coarsening Dynamics

The thermodynamic analysis suggests that the chromatin network acts much like rubber with entropic springs to hinder the coalescence of droplets. A similar mechanism could potentially impact the coarsening dynamics and the pathway leading to the formation of multiple droplets. To better understand the role of protein-chromatin interactions in the overall phase separation process, we analyzed the dynamical trajectories at the onset of cluster formation.

We first monitored the time evolution of the number of clusters formed along the dynamical trajectories shown in Fig. 2. The clusters were identified as high-density regions of protein particles across the entire nucleus using the DBSCAN (Density-Based Spatial Clustering of Applications with Noise) algorithm [61] (see Supplementary Materials). A typical trajectory is shown in Fig. 5A, and starts with zero clusters due to the random distribution of protein particles in the initial configuration. The sudden appearance of nine clusters at time ~ 14 min suggests that nucleation can occur at multiple sites almost simultaneously. As time proceeds, the droplets began to merge or evaporate, and the trajectory eventually stabilizes to the two-droplet state.

**FIG. 5.**
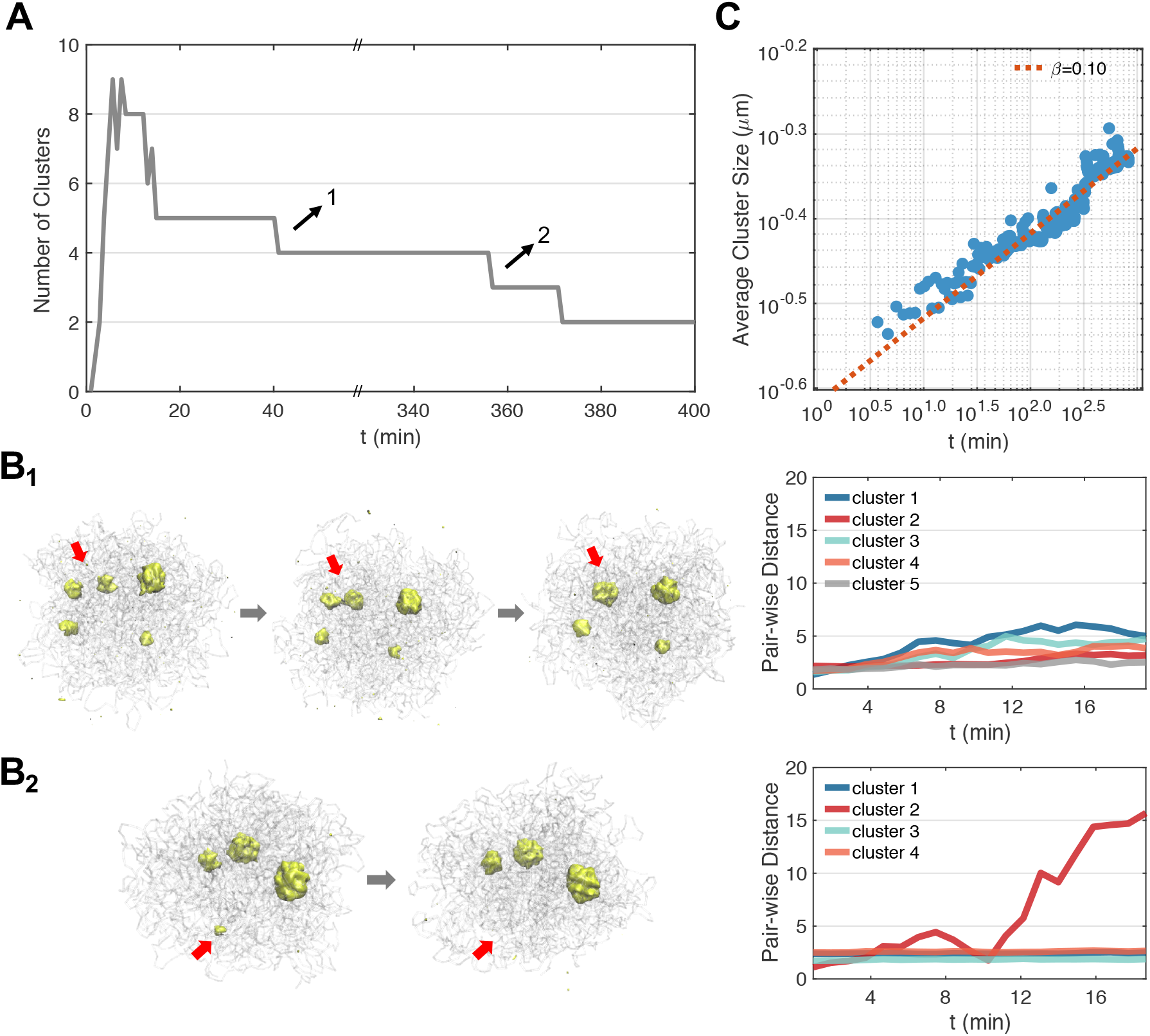
Coarsening Dynamics and pathways for phase separation with the chromatin network. (A) Time evolution of the number of clusters observed along a dynamical trajectory. (B) Detailed characterization of the two switching events that follow the Brownian motion-induced coalescence (B_1_) and the diffusion-limited Ostwald ripening (B_2_) path. The average pair-wise distance for each cluster remains relatively constant along the first path but increases significantly along the second one due to particle evaporation. (C) Power-law scaling of the average cluster size as a function of time.

We further recorded the average size of all clusters measured by their mean *R_g_*. The increase of *R_g_* as a function of the time follows a power-law scaling with *R_g_* ∝ *t^β^*. The exponent *β* = 0.1 differs from the phase separation process without the chromatin network. As shown in Fig. S9, for simulations performed without the network, the initial cluster number is smaller and the exponent for cluster size growth is close to the theoretical value of 1/3 [62, 63].

To provide insight into the appearance of the abnormal exponent *β*, we characterized the switching events that led to the decrease in cluster numbers. Specifically, we tracked the average pair-wise distance 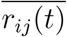 of protein particles within each cluster. In some cases, 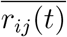 for all clusters remain relatively constant and the switching occurs through the Brownian motion-induced coalescence (BMC) pathway (Fig. 5B1). In other cases, we observe a significant increase in 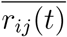 before the switching occurs (Fig. 5B2). This increase indicates an “evaporation” of the cluster following the diffusion-limited Ostwald ripening (DOR) path. We then identified all the switching events (in total 42) from the dynamical simulations and found that approximately 76% of them follow the BMC path. The rest of the switching events proceed via the DOR path and often involve smaller clusters to reduce the penalty associated with cluster disassembly (Fig. S10).

The dominance of the BMC pathway offers an explanation for the dramatic slowdown of the coarsening dynamics. In particular, the scaling exponent of 1*/*3 was predicted based on a normal diffusion model in which the distance between droplets scales linearly in time, i.e., *x*^2^(*t*) ∝ *Dt* [64]. However, as shown in Fig. S11, most of the clusters exhibit sub-diffusion and *x*^2^(*t*) ∝ *Dt*^1/2^. Sub-diffusive motion is indeed common inside the nucleus [65–67], and arises as a result of the elastic stress from the viscoelastic environment consisting of chromatin and nucleoplasm [67–70]. Assuming that the average size of droplets *r*(*t*) is proportional to their mean distance, we arrive at *r*(*t*) ∝ *t*^1/6^ for the observed abnormal diffusion. The exponent now is closer to the value shown in Fig. 5C.

We note that the abnormal diffusion and the slower coarsening kinetics are consistent with in vivo experimental results performed by Brangwynne and coworkers [19]. In particular, they revealed a coarsening exponent of ~ 0.12, which is close to the value shown in Fig. 5C.

### Protein-Chromatin Interactions Promote Cluster Nucleation

Our results indicate that, while protein-chromatin interactions increase the overall stability of the single-droplet state, they retard the coarsening kinetics by giving rise to sub-diffusion and entropic barriers. To more directly probe their impact on droplet coalescence, we recomputed the free energy profile at stronger (*ϵ* = 2.0 *k*_B_*T*) and weaker (*ϵ* = 1.6 *k*_B_*T*) protein-chromatin interactions. As shown in Fig. 6A, while the stability of the merged state varied significantly, the transition and the two-droplet state are much less affected. These simulations again support the entropic origin of the free energy barrier.

**FIG. 6.**
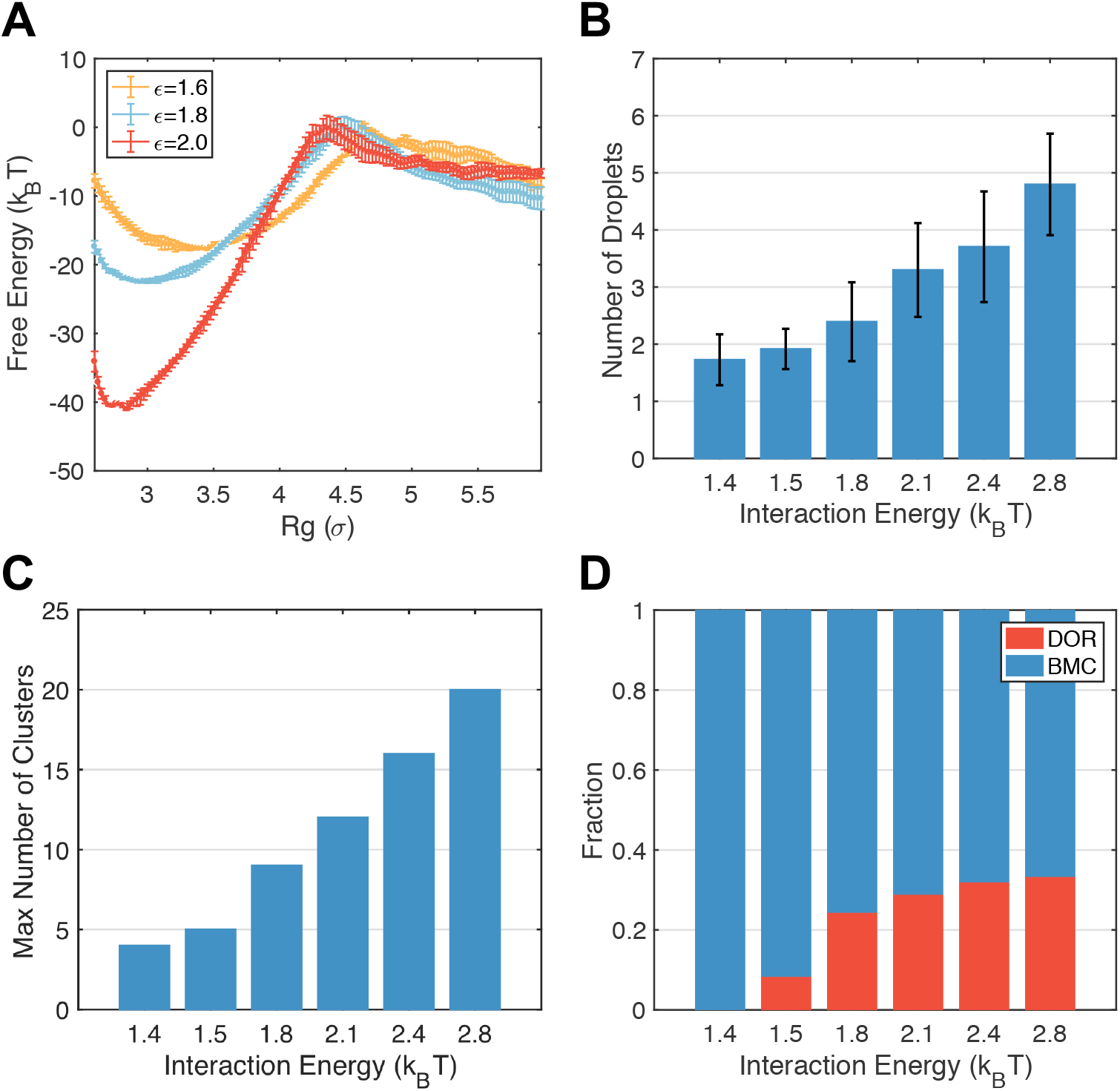
Protein-chromatin interactions promote cluster nucleation. (A) The free energy profiles of droplet coalescence at various protein-chromatin interactions. The result for *ϵ* = 1.8 is identical to the one presented in Fig. 3. Error bars were calculated as standard deviation of the mean. (B) Number of droplets formed at the end of dynamical simulations at various strength of protein-chromatin interactions. Error bars correspond to standard deviations of results from independent trajectories. (C) The maximum number of clusters observed in dynamical trajectories performed at various protein-chromatin interactions. (D) The fraction of cluster switching events following the Brownian motion-induced coalescence (BMC, blue) and the diffusion-limited Ostwald ripening (DOR, red) at various protein-chromatin interactions.

We performed additional long-timescale simulations following the same protocol as those shown in Fig. 2. Starting from random protein configurations, we varied protein-chromatin interactions from 1.4 *k*_B_*T* to 2.8 *k*_B_*T* and carried out eight independent six-million-step long simulations for each interaction strength. We then computed the number of droplets formed at the end of these simulations. As shown in Fig. 6B, the number of droplets increases with the interaction strength. Stronger protein-chromatin interactions facilitate the nucleation of protein clusters on the chromatin, as can be seen from the increase of cluster numbers at the onset of phase separation (Fig. 6C). Cluster coarsening, again, primarily follow the BMC pathway, though the ratio of DOR increases as well due to the instability of the nucleated clusters with smaller number of protein particles (Fig. 6D). Since merging of these clusters along the BMC pathway is hindered by entropic barriers, more nucleation naturally leads to increased droplet numbers at the end of the simulations. These results directly support the role of the chromatin network in hindering droplet coalescence.

## DISCUSSION

While liquid-liquid phase separation has emerged as the leading mechanism for nuclear body formation, the impact of the surrounding nuclear environment on nuclear bodies is less known. We studied the role of protein-chromatin interactions for phase separation inside the nucleus. Our simulations used the diploid genome model parameterized from Hi-C data to provide an accurate description of the chromatin network. We found that protein-chromatin interactions facilitate the nucleation of condensates and hinder their coalescence, stabilizing the multiple-droplet state.

Many prior studies support the importance of protein-chromatin interactions in regulating nuclear body numbers. For example, while two nucleoli are often seen in normal cells, the number increases significantly upon tumorigenesis [54]. Enhanced protein-chromatin interactions resulting from elevated rDNA transcription may well underlie this increase. Paraspeckle numbers have been shown to correlate with the nucleus size [20]. Such a correlation could arise as the entropic barrier shown in Fig. 3 becomes more evident at larger nuclei to further suppress droplet coalescence. Finally, reducing protein chromatin interactions upon inhibiting transcription was found to facilitate the coalescence of speckles [71], a result that’s consistent with the loss of the free energy barrier.

It’s worth mentioning that in addition to protein chromatin interactions, many other factors play important roles in stabilizing the nuclear bodies. In particular, since phase separation only occurs when the concentration exceeds a certain threshold, changes in cell volume that alter protein concentration could impact nucleoli size and number [72]. In addition, many nuclear bodies are involved in chemical reactions such as splicing and transcription. The active production of new molecules and the post-translational modification of proteins could introduce non-equilibrium mechanisms that may contribute to the formation of multiple droplets [73]. Finally, the nuclear envelope could be another factor, as evidenced by the fusion of nucleoli in senescence cells that lost lamin-chromatin interactions [74, 75]. Whether the effect is propagated via the chromatin network or through direct interactions with nucleoli remains to be shown. Accounting for these additional factors could provide a more comprehensive understanding of in vivo phase separation and nuclear body formation.

## METHODS

### Setup of the Nucleus Model

In addition to the 46 chromosomes that follow the same setup and interactions as in Ref. [27], we introduced a total of 500 particles to represent nucleolar proteins. The number of particles was chosen to reproduce a protein concentration of 1 *μ*M [76] in nucleoli of approximately 1 *μ*m in diameter. The size of the protein particles *σ*_p_ was further estimated to be 0.5*σ* assuming a tight packing when forming the nucleoli (see Supplementary Materials). We note that from our estimation, it is clear that these particles do not represent a single specific protein but correspond to molecular aggregates of protein and RNA.

In addition to their attractive self-interactions, protein particles bind with chromatin via specific and non-specific interactions. All three types of interactions were modeled with the cut and shifted Lennard-Jones potential

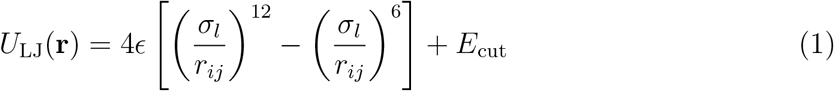

for *r_ij_* < *r_c_* and zero otherwise, where *r_c_* = 2.0*σ*. 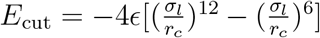 *ϵ* = 1.8, 1.8 and *k*_B_*T*, and *σ_l_* = 0.5, 0.75 and 0.75*σ* for protein-protein, protein-NAD and protein-nonNAD interactions. NADs were identified using the high-resolution sequencing data generated from Ref. [33]. We processed the raw nucleolar-to-genomic ratios to generate signal data at the 1 Mb resolution. Only genomic loci with signals higher than a threshold value (15.0) were labeled as NADs and homologous chromosomes share identical NADs.

### Details for Molecular Dynamics Simulations

We used the molecular dynamics package LAMMPS [77] to perform polymer simulations in reduced units. Constant-temperature (*T* = 1.0 in reduced unit) simulations were carried out via the Langevin dynamics with a damping coefficient *γ* = 10.0*τ_B_* and a simulation time step of *dt* = 0.008*τ_B_*, where *τ_B_* is the Brownian time unit. As shown in the Supplementary Materials, *τ_B_* can be mapped to the real-time unit of 0.028 s using a nucleoplasmic viscosity of 10^−2^ Pa·s (see Supplementary Materials).

### Simulation Details for Free Energy Calculations

We computed the free energy profile for coalescence using umbrella sampling and temperature replica exchange with 16 umbrella windows [78, 79]. There simulations were initialized from a typical two-droplet configuration and lasted for forty million steps. Indices of protein particles in the two droplets were identified using the DBSCAN algorithm (see Supplementary Materials). We defined the collective variable for these particles as 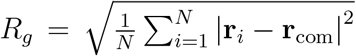. **r**_*i*_ is the coordinate of each protein particle and **r**_com_ is the center of mass. A harmonic potential 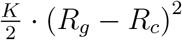 was introduced in each umbrella window to facilitate the sampling of configurations at targeted distances. We chose the center of these windows, *R_c_*, to be evenly spaced between 2.5 and 6.0 with an increment of 0.25, except for the first window whose *R_c_* = 2.0. The spring constant *K* was chosen as follows:

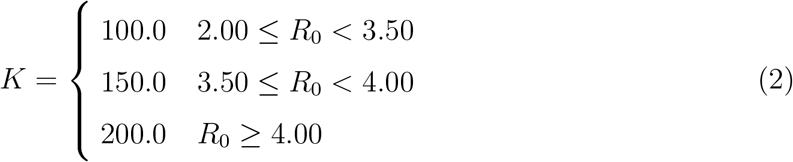

Eight replicas were used within each umbrella window with temperatures ranging from 1.00 to 1.14 with 0.02 increments. Exchanges between these replicas were attempted at every 100 time steps.

## ACKNOWLEDGEMENT

We thank Dr. Xingcheng Lin, Dr. Kartik Kamat and Dr. Xinqiang Ding for helpful discussions. This work was supported by the National Institutes of Health (Grant R35GM133580).

